# *Bacillus subtilis* remains translationally active after CRISPRi-mediated replication initiation arrest

**DOI:** 10.1101/2022.09.20.508806

**Authors:** Vanessa Muñoz-Gutierrez, Fabián A. Cornejo, Katja Schmidt, Christian K. Frese, Manuel Halte, Marc Erhardt, Alexander K.W. Elsholz, Kürşad Turgay, Emmanuelle Charpentier

**Affiliations:** Max Planck Unit for the Science of Pathogens, D-10117 Berlin, Germany; Institute of Microbiology, Leibniz Universität Hannover, D-30419 Hannover, Germany; Humboldt-Universität zu Berlin, Institute for Biology – Bacterial Physiology, Philippstr. 13, D-10115 Berlin, Germany; Institute for Biology, Humboldt-Universität zu Berlin, D-10115 Berlin, Germany

**Keywords:** *Bacillus subtilis*, replication, *oriC*, DnaA boxes, CRISPRi

## Abstract

Initiation of bacterial DNA replication takes place at the origin of replication, a region characterized by the presence of multiple DnaA boxes that serve as the binding sites for the master initiator protein DnaA. The absence or failure of DNA replication can result in bacterial cell growth arrest or death. Here, we aimed to uncover the physiological and molecular consequences of stopping replication in the model bacterium *Bacillus subtilis*. For this purpose, DNA replication was blocked using a CRISPRi approach specifically targeting DnaA boxes 6 and 7, which are essential for replication initiation. We characterized the phenotype of these cells and analyzed the overall changes in the proteome using quantitative mass spectrometry. Cells with arrested replication were elongating and not dividing but showed no evidence of DNA damage response. Moreover, these cells did not cease translation over time. This study sets the ground for future research on non-replicating but translationally active *B. subtilis*, which might be a valuable tool for biotechnological applications.

**Importance:** Even though bacteria are constantly replicating under laboratory conditions, natural environments expose them to various stresses like lack of nutrients, high salinity, and pH changes, which can keep them in non-replicating states. Non-replicating states can allow bacteria to become less sensitive or tolerant to antibiotics (persisters), remain inactive in specific niches for an extended period (dormancy), and adapt to some hostile ecosystems. Non-replicating states have been studied due to the possibility of repurposing energy to produce additional metabolites or proteins. Using CRISPRi targeting bacterial replication initiation sequences, we successfully arrested the replication of *B. subtilis*. We observed that non-replicating cells continued growing but not dividing, and the initial arrest did not induce global stress conditions such as SOS or stringent response. Notably, these cells continued their metabolic activity and translation. This study provides comprehensive insights into the physiological response of replication initiation blockage in *B. subtilis*.

## Introduction

DNA replication aims to duplicate the genomes of dividing cells to contribute a new identical DNA copy to each daughter cell. Replication is therefore connected to cell growth, chromosome segregation, and cell division. In bacteria, replication initiates at a specific part of the chromosome, the origin of replication (*oriC*). This region contains several DNA sequences called DnaA boxes that serve as binding sites for the master initiator protein DnaA. DnaA binds to several boxes at the *oriC*, causing the unwinding of an AT-rich region within the *oriC* and facilitating the recruitment of the replication machinery at both strands (1–4). The absence or failure of DNA replication can lead to bacterial growth arrest or cell death. Although bacteria can rapidly divide and replicate under laboratory conditions, most bacteria in their natural habitat encounter multiple stresses, such as nutrient scarcity, that hinder the capacity to duplicate. One example of the non-replicative states in bacteria is persister cells, which constitute a small percentage of the bacterial population that stops growing and is, therefore, less sensitive to antibiotics (5). Another example is the “oligotrophic growth state”, in which bacteria can survive for multiple days in pure water while maintaining basic metabolic activities (6).

The physiology of non-growing bacteria has been studied due to its relevance in industrial and clinical contexts (7). The main advantage of using these cells in industry biotechnology is that the carbon and energy spent for biomass formation can be redirected to produce compounds of interest (8, 9). Notably, metabolically active states in non-growing bacteria have previously been reported, with some displaying enhanced protein production (10–13). The methods to induce these non-replicating states include, for example: **i)** the use of toxins from toxin-antitoxin systems to induce a persister-like phenotype (14, 15), **ii)** exploiting repressible promoters to downregulate the RNA polymerase (10), **iii)** the elimination of chromosomal content with endonucleases such as I-CeuI (11), **iv)** the removal of the *oriC* with serine integrases (16), **v)** mimicking nutrient depletion, the addition of growth inhibitors, and the use of CRISPRi to knockdown genes related to growth or replication (8, 9, 17, 18). In this study, we constructed a strain with a chromosomally integrated CRISPRi system specifically targeting DnaA boxes 6 and 7, which have been previously reported to be the most important to promote DNA unwinding during replication initiation in *B. subtilis* (19). We characterized the phenotype upon CRISPRi induction by evaluating cell morphology, membrane integrity, replication status, and SOS response. The time-resolved quantitative proteome profile of this strain reveals a significant change in the abundance of ribosomal proteins; however, SOS response proteins are not significantly elevated. This indicates that these cells stop replicating but continue growing and translating. Overall, our results shed light on the fundamental role of replication initiation in *B. subtilis* and provide a foundation for subsequent biotechnological studies of this bacterium as a producer strain.

## Results

### An inducible CRISPRi system to block replication initiation

A recent study showed that the DnaA boxes 6 and 7 on the *oriC* of *B. subtilis* are crucial for DNA unwinding and replication initiation (19). Thus, we hypothesized that initiation could be blocked by sterically hindering the binding of DnaA to these boxes using a xylose-inducible dCas9 (18) (Fig. 1A). As a control, we also selected sgRNAs targeting boxes 1 – 2 and 3 – 4 (sgRNA^box1-2^ and sgRNA^box3-4^) (Fig. 1B). This system allows the conditional expression of dCas9 when xylose is present in the medium but also tight repression of it when glucose is present.

**Figure 1:**
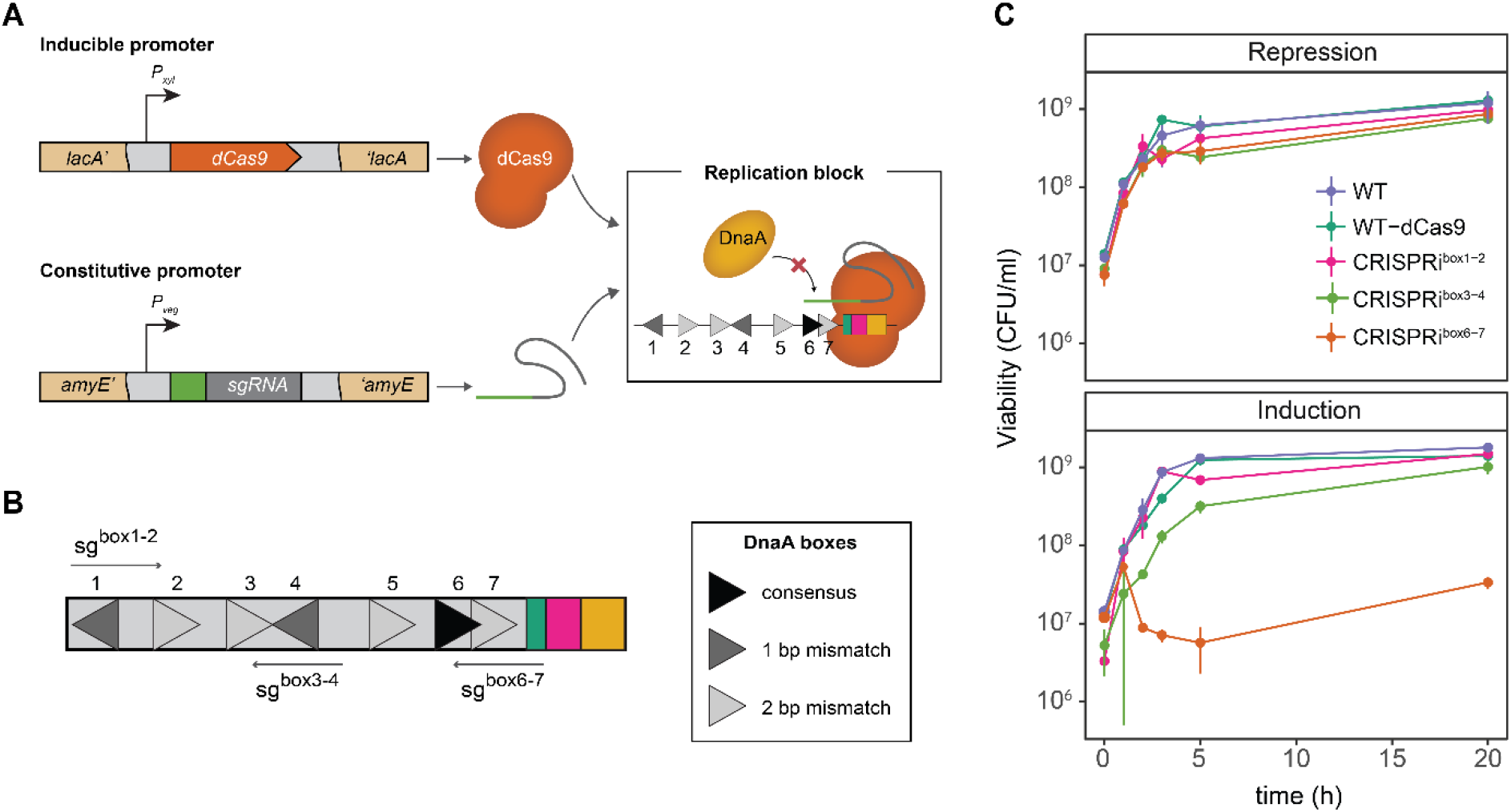
CRISPRi targeting the DnaA boxes 6-7 inhibit cell proliferation. (A) *B. subtilis* carrying a xylose-inducible dCas9 (orange) is directed to specific DNA targets by constitutively expressed sgRNAs (green) under the P_*veg*_ promoter control. The dCas9-sgRNA complex blocks the binding of DnaA (yellow) to the DnaA boxes (triangles). *dcas9* was stably integrated into the *lacA* locus, and the sgRNAs were integrated into the *amyE* locus. (B) DnaA boxes from *B. subtilis* and the selected sgRNA targets (left side). Other elements of the *oriC* are represented: DnaA trios (green), DnaD binding sites (fucsia) and the AT-rich region (yellow). (C) CRISPRi^box6-7^ does not resume growth when induced at the early exponential growth phase. Cell counts of cells under repression (glucose) or induction (xylose) conditions. Data shown is the mean of 3 biological replicates and the error bars represent the SD.

To assess the block of replication initiation, we evaluated the growth of cells in the early logarithmic phase under induction or repression of dCas9. Cells carrying only the dCas9 integration grew similarly in inducing and repressing conditions, meaning that dCas9 expression alone does not alter the capacity of bacteria to proliferate (Fig. 1C). Repressed cultures of the sgRNA^box1-2^ and sgRNA^box3-4^ had slight growth differences compared to the control (Fig. 1C). Inducing dCas9 in the strain containing the sgRNA^box6-7^ leads to a significant decrease in CFU/ml after 3 h. On the contrary, we did not observe growth inhibition when the CRISPRi system targeted the other adjacent boxes (Fig. 1C). This indicates that the observed effect results from a competition between DnaA and dCas9 for binding to the DnaA boxes 6 and 7. This specific block by the dCas9-sgRNA^box6-7^ complex could prevent DnaA-filament formation and the subsequent associated replication-bubble opening that starts replication. These observations also confirm the high specificity of DnaA binding to certain boxes, specifically DnaA boxes 6 and 7, as an initial start of replication initiation, as previously reported (19).

### CRISPRi system inhibits the initiation of replication, leading to replisome disassembly

To verify the effect of CRISPRi blocking replication initiation at single cell level, we followed the replisome localization by generating a translational fluorescent reporter gene fusion to the gene encoding the beta clamp of the polymerase protein, DnaN, at its endogenous genomic locus. DnaN colocalizes with the replisome making it an excellent reporter to evaluate the progression of DNA replication in the cell (20). The GFP-DnaN fusion protein forms bright foci that assemble and disassemble inside the cells, signaling the start and completion of DNA replication, respectively. Previous studies have shown that no DnaN foci form in the absence of replication, while actively replicating cells contain between zero to four DnaN foci (20, 21).

Dynamic localization of GFP-DnaN was evaluated under xylose conditions (Fig. 2A). The wild-type (WT) strain displayed bright foci, indicating that the replication forks are assembled, and the cells are actively replicating. Remarkably, in strain CRISPRi^box6-7^, foci diffusion is already observed after 2 hours of xylose induction, suggesting that replication is no longer initiated in these cells.

**Figure 2.**
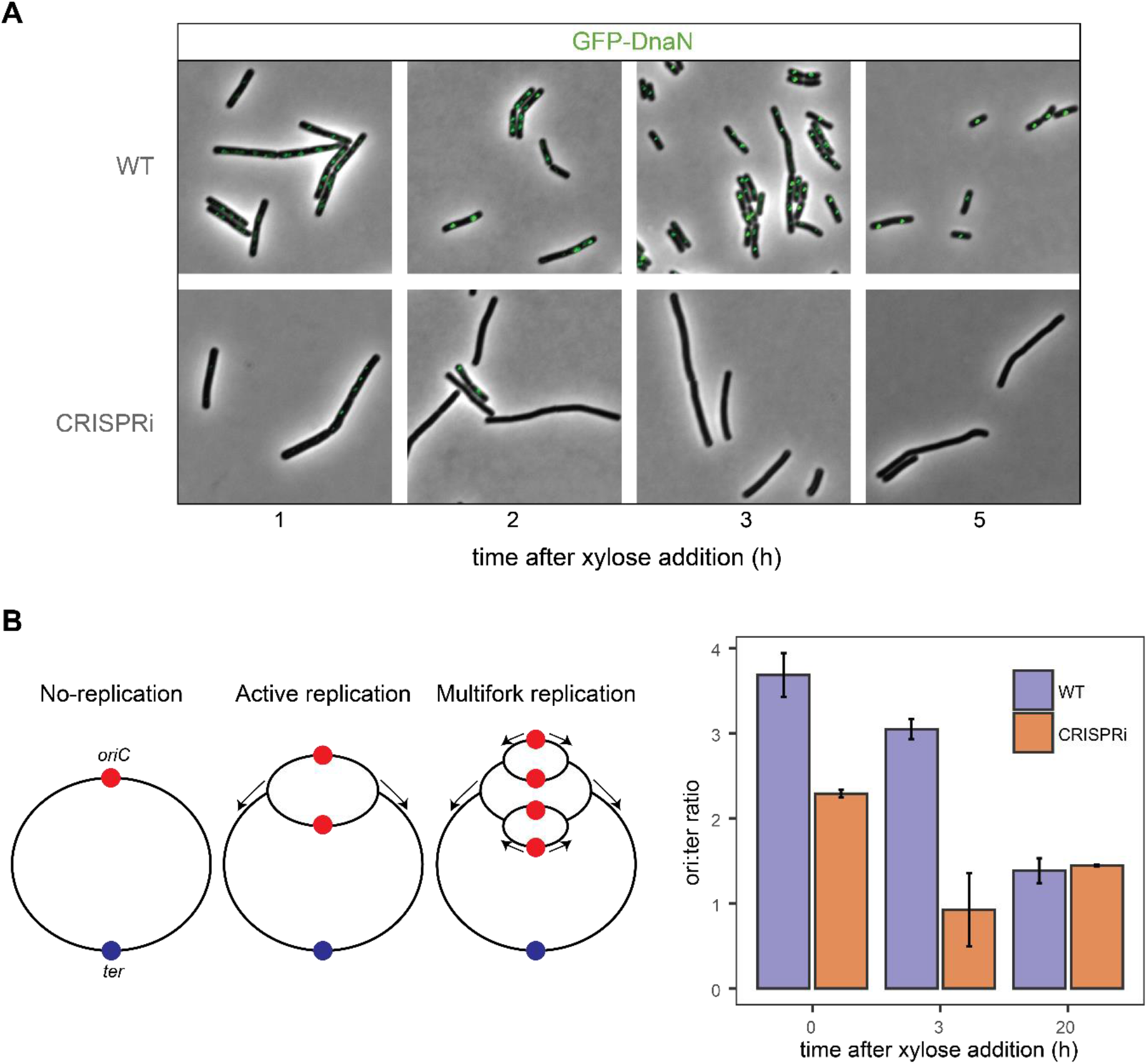
Replication is inhibited in the CRISPRi strain. (A) Epifluorescence microscopy of *B. subtilis* cells expressing GFP-DnaN undergoing CRISPRi replication arrest. Strains were grown in LB + 1% glucose until OD600nm 0.1, washed and resuspended in LB + xylose 1%. Cells were immediately subjected to microscopic analysis at 1, 2, 3 and 5 h after xylose addition. (B) Left panel: Explanation of replication states in bacteria. In nutrient-rich conditions, chromosomes undergo multifork replication and have more than one *oriC* (red) per cell; therefore, their *ori* to *ter* (blue) ratio is higher than in non-replicating conditions. Right panel: ori:ter ratios as determined by RT-qPCR for WT and CRISPRi cells. Data points in (B) represent the means of three independent replicates; error bars indicate standard deviations of the mean.

Another method to demonstrate that replication initiation has stopped is analyzing the frequency of the origin and terminus region (ori-ter ratio). Cells that are actively replicating have a ratio above 1, while cells containing only one copy of the chromosome, like stationary phase cells or in a non-replicating state, are expected to have a ratio closer to 1 (Fig. 2B). We determined the ori-ter ratios using qPCR (Fig. 2B, Table S3). We observed that blocking the DnaA boxes 6 and 7 resulted in a decreased DNA replication initiation rate after 3 hours of induction with an ori:ter ratio of 0.88 vs 2.89 for WT cells. This demonstrates that replication is blocked by the designed system. These observations are consistent with the inhibition of DNA replication initiation and were confirmed by the observed replisome disassembly with the GFP-DnaN reporter.

### No evidence of DNA damage response under CRISPRi-mediated replication arrest

Bacteria can sense and respond to DNA stress through the widely conserved SOS response (reviewed in (22, 23)). The SOS response initiates when single-stranded DNA accumulates following DNA damage or replication blocks. One of the possible consequences of inducing replication arrest in a rich medium is the occurrence of replication-transcription conflicts at stalled replication forks resulting in DNA damage (24). In *B. subtilis*, the protein involved in inhibiting cell division during the SOS response is YneA (25), whose expression is repressed by LexA, the transcriptional repressor of the SOS regulon (26).

To test whether the replication-arrested cells were undergoing DNA damage response (DDR), we added an ectopic integration of *gfp* under the control of the *yneA* promoter in the CRISPRi and WT background strains. Since DNA damage leads to de-repression of genes controlled by LexA such as *yneA* (27), we used GFP expression as an indicator of SOS response. For this, the GFP fluorescence of the cells was observed at various timepoints after xylose addition. As a positive control, we performed the same experiment but exposed the cells to mitomycin C, a DNA-damaging agent which creates interstrand cross-links, induces the SOS response and results in cell elongation (25).

Both WT and CRISPRi strains showed no signs of DDR using the P_*yneA*_-GFP reporter compared to the mitomycin DDR response in WT cells (Fig. 3A). This observation indicates that the SOS response was not induced during CRISPRi-mediated replication arrest.

**Figure 3.**
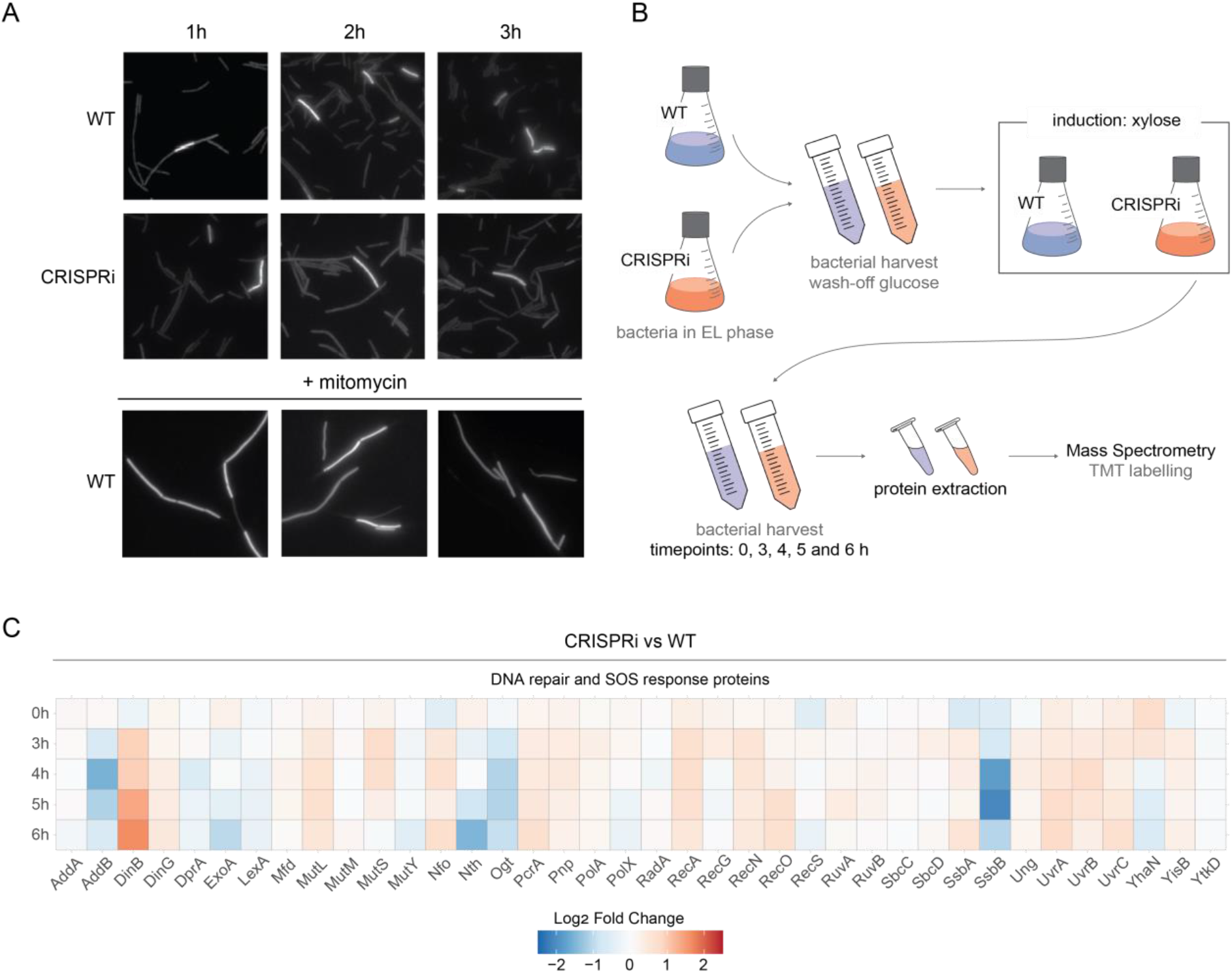
No evidence of DNA damage under replication arrest. (A) WT and CRISPRi strains containing the P_*yneA*_-*gfp* ectopic integrations and grown LB with 1% xylose. Images are from aliquots taken from cultures constantly growing at 37°C. Cells were subjected to microscopic analysis at 1, 2 and 3 h after xylose addition. (B) An overview of the experimental setup to collect the samples for MS analysis. WT and CRISPRi cells were harvested at different timepoints after xylose induction, and their proteins were extracted and quantified by Tandem Mass Tag (TMT) mass spectrometry. EL: Early logarithmic phase. (C) Heatmap based on Fold Change values of selected proteins. DNA repair and SOS response-associated proteins are retrieved from SubtiWiki. Data selected presents mean of 3 biological replicates.

To explore the response of *B. subtilis* to CRISPRi-mediated replication arrest over time, we analyzed the overall proteome for differences between CRISPRi and WT samples using quantitative mass spectrometry (Fig. 2B).

We observed almost no difference in the protein abundance profiles between replicating cells (WT) and non-replicating cells (CRISPRi) at timepoint 0 h and clear changes in differentially expressed proteins over the course of the experiments (Fig. S1). Additionally, we observed that the DNA repair and SOS response proteins of CRISPRi vs. WT did not show clear differences except for the nuclease inhibitor DinB, which is differentially more abundant under replication arrest (Fig. 3C). These two observations add further evidence that there is no significant DNA damage response in the replication-arrested cells.

### CRISPRi-mediated replication arrest enhances protein expression

To obtain a global view of the change in cellular processes upon replication arrest, we analyzed the enrichment of the Gene Ontology (GO) terms within these cells (Fig. 4A and Fig. S2). Here, we observed that after 5 and 6 h of replication arrest, the ribosomal proteins and other translation-related proteins were more abundant. Our analysis indicated no enrichment of SigB-regulated general stress response proteins (Fig. S3).

**Figure 4.**
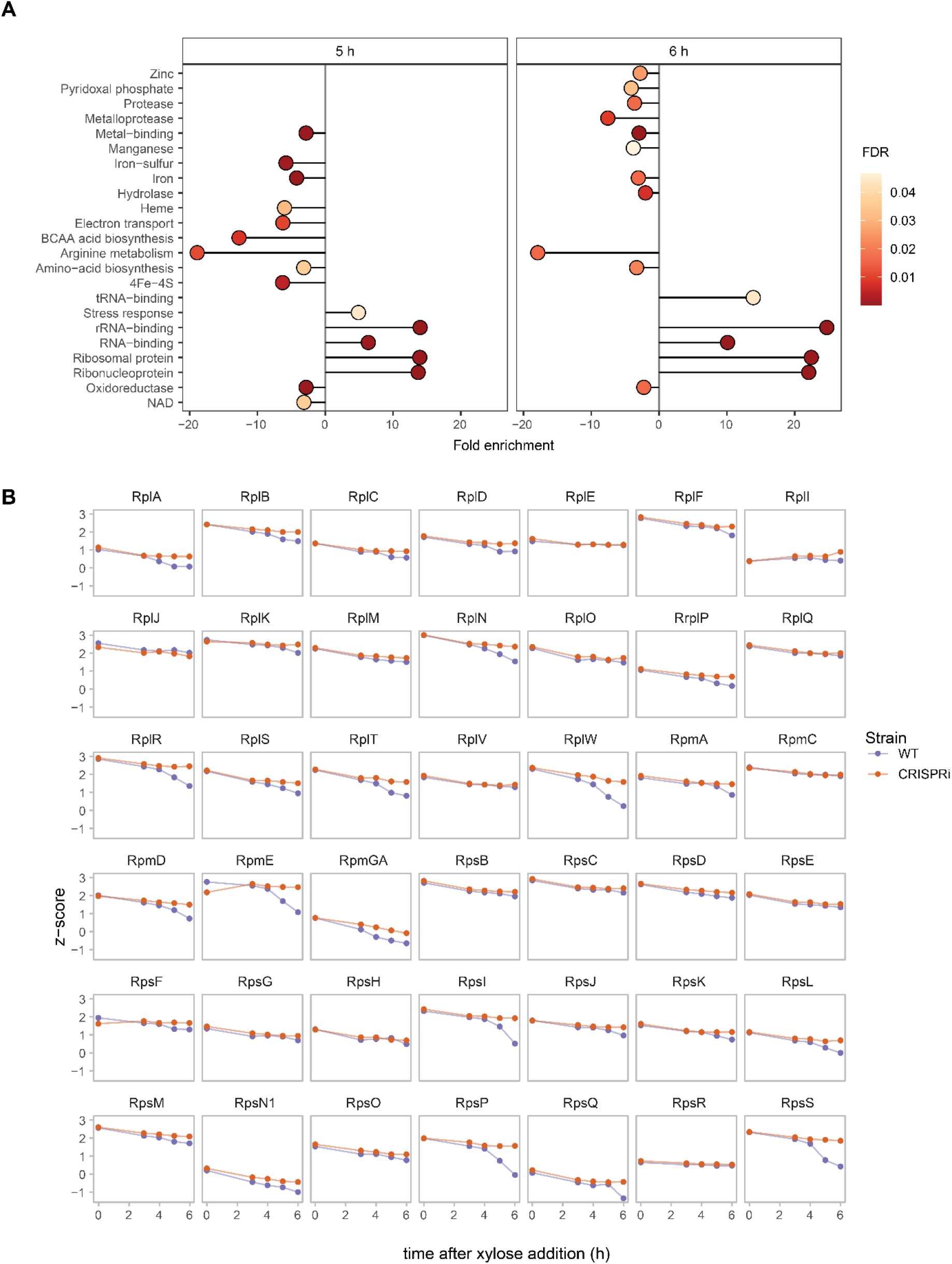
Proteomic characterization shows translation-related proteins are more abundant in replication-arrested cells after 5 and 6 h of xylose induction. (A) GO enrichment analysis of differentially expressed genes in CRISPRi and WT strains. Proteins whose abundance was significantly changed were analyzed. The enriched terms after 5 and 6 h of xylose induction are shown. Dot color relates to False discovery rate (FDR) values for each process. Profile plots of z-score ribosomal proteins retrieved from Subtiwiki. Purple line indicates WT and yellow CRISPRi strain.

Enhanced translation upon replication arrest is a possible mechanism to increase protein and metabolite production and has been studied in *E. coli* (8, 9, 28). To observe whether the levels of translation-related proteins were increased upon replication arrest in *B. subtilis*, we made individual profile plots of translation-related proteins as a function of the z-score (Fig. 4B). We observed that the level of most ribosomal proteins in the CRISPRi strain remains constant over time. In contrast, protein levels in the WT cells decreased (Fig. 4B). Our interpretation of this result is that as WT cells enter the stationary phase, CRISPRi cells appear to be in an extended exponential phase from the moment the replication block with dCas9 happens.

Next, we wanted to assess if the ribosomal protein abundance was related to an increased translation. Therefore, we generated an ectopic integration of *gfp* under an IPTG inducible promoter (P_*hyperspank*_) in the WT and CRISPRi background strains as a reporter for protein translation. We then measured GFP fluorescence in a time-lapse experiment after xylose and IPTG addition (Fig 5A). We note that the cytoplasmic GFP signal allowed us as well to observe the general morphology of the cells. WT cells showed clear cell division and septum formation. In contrast, CRISPRi cells showed an elongated phenotype without an apparent septum (Fig. 5A). We next quantified the fluorescence intensity of individual cells from the timelapse microscopy experiment. We found that after 3 hours of xylose addition, CRISPRi cells displayed a significantly enhanced GFP intensity per cell compared to the WT. These results at the proteomic and single-cell levels indicate that replication arrest affects the overall translation process. However, the observed increase in GFP expression might also be caused by its stabilization.

**Figure 5.**
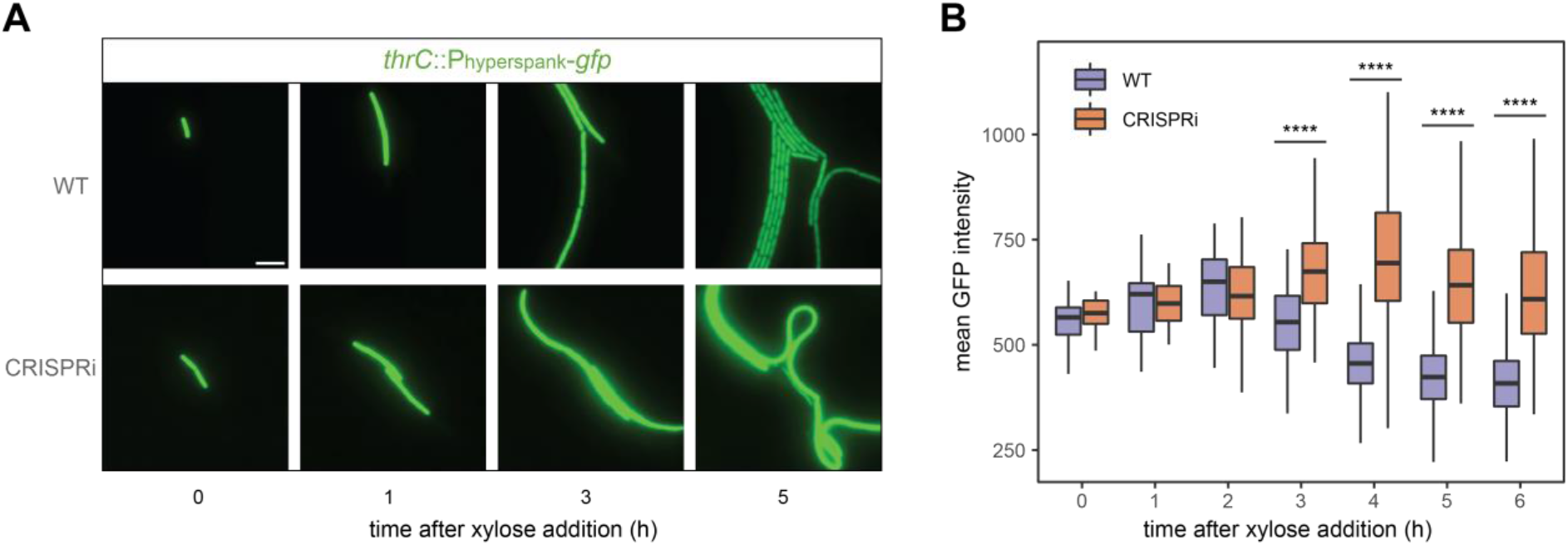
Chromosomal GFP expression is higher after replication arrest. (A) Snapshots taken from a time-lapse microscopy experiment. An ectopic integration of *gfp* under an IPTG inducible promoter was analyzed in the WT and CRISPRi background strains. Strains were grown in the presence of both xylose and IPTG. CRISPRi cells showed an increase in fluorescence. Representative pictures are shown. (B) Mean GFP intensity per cell upon xylose induction in WT and CRISPRi cells in a time-lapse experiment (as panel A, Video S1). The timelapse was performed in 3 biological replicates, each replicate followed 4 areas per strain. Each area contained at least 3 cells. Two-tailed Welch tests were performed for CRISPRi against WT as a control group and asterisk represent p-values. *: P ≤ 0.05, **: P ≤ 0.01, ***: P ≤ 0.001, ****: P ≤ 0.0001.

### CRISPRi-mediated replication arrest causes cell elongation

From the time-lapse experiment, we next quantified the cellular morphology after inducing replication arrest. Overall, cells are larger during replication arrest in the CRISPRi cells. These differences became significant already after 1 hour of xylose induction and became even more significant after 2 hours of induction and throughout the whole experiment (Fig. 6A).

**Figure 6.**
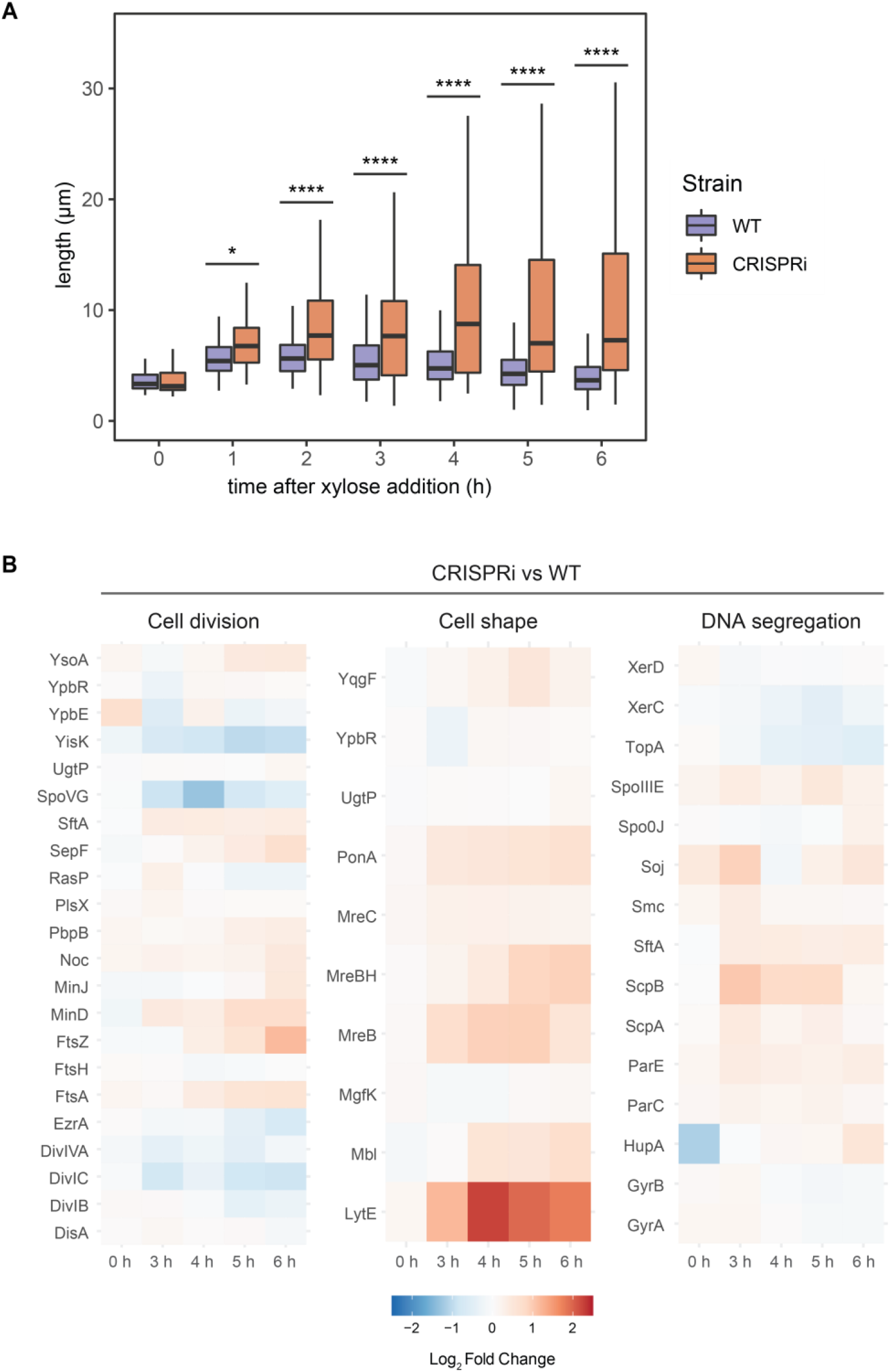
Uncontrolled cell elongation under replication arrest. (A) Quantitative analysis of cell length from time-lapse data (Fig. 5A and Video S1). Cell length increases upon addition of xylose in CRISPRi cells. The timelapse was performed in 3 biological replicates, each replicate followed 4 areas per strain. Each area contained at least 3 cells. Two-tailed Welch tests were performed for CRISPRi against WT as a control group and asterisk represent p-values. *: P ≤ 0.05, **: P ≤ 0.01, ***: P ≤ 0.001, ****: P ≤ 0.0001. (B) Heatmap based on Fold Change values of proteins related to cell division, cell shape and DNA segregation. All proteins annotations are retrieved from SubtiWiki.

To pinpoint specific proteins that can be enriched in the CRISPRi cells and explain the elongated phenotype, we analyzed the protein abundance in our proteomic experiment for other related processes like cell division, cell shape, and DNA segregation (Fig. 6B). We observed that most proteins have similar cellular levels compared to the WT. However, we found three related proteins: LytE, MreB, and MreBH, to be differentially more abundant in CRISPRi (Fig. 6B). LytE is a cell wall hydrolase involved in cell elongation and separation (29), whose activity requires functional MreB and MreBH (30, 31). This observation suggested that uncontrolled cell elongation might lead to cell lysis, which we observed after 7 hours of xylose induction (Video S1).

Taken together, these results show that CRISPRi mediated replication initiation arrest is specific for the DnaA boxes 6 and 7. Moreover, this system stop the process of cell division, while continuing the processes of cell translation and elongation. Additionally, replication arrested cells do not undergo global stress response regulation.

## Discussion

In this work, we employed a rational design using CRISPRi to block replication initiation in *B. subtilis*. This block hinders the capacity of DnaA to bind specific DnaA boxes, thereby impeding replication initiation but not interfering with DnaA expression and its function as a transcriptional regulator (32). We found that targeting DnaA boxes 6 and 7 was more efficient than targeting other DnaA boxes within the *oriC* (Fig. 1C). This finding aligns with previous reports targeting DnaA boxes in *B. subtilis* (19). In contrast, targeting any DnaA box within the *oriC* in *E. coli* produces a non-replicating phenotype (17). Based on these observations, we confirm that specific DnaA boxes within the *oriC* in *B. subtilis* have a specialized function to trigger replication initiation. However, it remains elusive whether such functional hierarchy in the DnaA boxes in *B. subtilis* is observed in other bacteria. We expect this to be the case given the conserved *oriC* architecture in *B. subtilis* and its similarities with other bacteria (33). It is also interesting to know to what extent the location of *oriC* in *E. coli*, which is 40 kb away from the DnaA and DnaN genes, might affect the functional hierarchy of the DnaA boxes (33).

CRISPRi-mediated replication arrested cells were characterized by employing cell biological assays in combination with proteomics approaches. We found that CRISPRi-mediated block at the DnaA boxes 6 and 7 leads to a shutdown of the critical cell cycle processes of DNA replication (Fig. 2) and cell division (Fig. 5A) while growth (Fig. 5A and Fig. 6A) and translation continue (Fig. 4A and Fig. 5B). The phenotype observed under replication arrest, where cells stop dividing and form filaments is characteristic of cells under the SOS response and is mediated by YneA (26). However, profile plots of SOS-response-associated proteins indicate that this stress response is not active (Fig. 3C). Moreover, the translational fusion of the *yneA* promoter to GFP to detect the cellular SOS response did not show activity (Fig. 3A and Fig. 3C).

Since the observed phenotype was not caused by the SOS-response, we think this could indicate that the stop in cell division is SOS response independent. SOS-independent inhibition of cell division has been reported in several microorganisms (34–36). Likewise, DnaA binds specifically to the promoter regions of several genes, including *ftsL* (32). FtsL is an unstable protein and is essential for cell division (34). A decrease in *ftsL* mRNA levels quickly causes a decrease in FtsL protein and inhibits cell division (25, 37). Future work can pinpoint at the specific response that halts division by following the promoter activity or the expression of those genes in non-replicating cells.

Proteomic analysis of the non-replicating cells revealed that proteins involved in translation (rRNA-binding, ribosomal proteins, tRNA-binding and nucleoproteins) are more abundant after 5 and 6 hour of xylose addition compared to WT (Fig. 4A). Studies in several bacteria have suggested that protein synthesis continues during growth arrest and that maintaining functional translation machinery may be required for their viability (10–12, 38). Additionally, several studies in *E. coli* demonstrated enhanced protein and biochemical production of proteins and metabolites during replication arrest (8, 9, 28). We further confirmed that non-replicating cells maintained a higher rate of protein expression than WT cells (Fig. 5).

While the effect of protein expression is enhanced as described in other organisms, replication arrested *B. subtilis* cells started to lyse after 7 hours of induction (Video S1) probably because of the uncontrolled elongation of the cells. Cell elongation has been previously reported in some temperature-sensitive mutants for replications initiation in *B. subtilis* (39–41) and CRISPRi-mediated replication arrest in *E. coli* (17). However, these studies did not address if the uncontrolled elongation results in cell lysis. These elongation effects may be bypassed by making the process reversible before cell lysis (on/off switch of replication arrest) using temperature switches (42), degradation tags (17), light-activated and sensitive variants (42–44), or split variants (45).

For future biotech applications of these replication inhibiting CRISPRi cells as a microbial cell factory, it is important to effectively remove the need to add xylose in the media to induce dCas9 and eliminate the cell elongation constraints. To address the first, the expression of dCas9 can be driven by self-inducing promoters (46–48) that remove the dependence on chemical inducers, which are not economical at industrial scales. For the second, keeping control of the cell shape and membrane homeostasis may extend the lifespan of the bacteria and avoid anticipated cell lysis for industrial applications (24). One way to address this issue is to knockout elongation proteins that are being overexpressed such as LytE (Fig. 6B). However, deletion of cell wall hydrolases like LytE might not have an effect due to their redundancy, as single knockouts present no phenotype. In a multiple knockout approach of *B. subtilis* cell wall hydrolases (42 in total), researchers were able to remove all but two of these genes in a single strain, *lytE* and *cwlO*, which were shown to be synthetically lethal (49). Their results indicate that the only essential function of cell wall hydrolases in *B. subtilis* is to enable cell growth by expanding the wall and that LytE or CwlO alone is sufficient for this function. Therefore, we suggest building a library to screen for targets that inhibit cell replication and growth while allowing for continued protein production. Similar approaches have been performed in *E. coli* (9).

The current system also has some limitations. For instance, comparing the non-replicating CRISPRi with the replicating WT cells is challenging. Over time, replicating cells increase cell density, so their nutrient availability is exhausted faster than non-replicating cells. Hence, up- or downregulation of specific metabolic pathways in replicating cells could also be caused by nutrient limitation and the entry into stationary phase. These concerns about a better-suited comparison of the two different cellular states can be addressed in future work by using either a chemostat or microfluidic approaches that can maintain the cells in similar growth states.

Finally, this system could become a source for producing many toxins and antibiotics that target the DNA replication machinery. In conclusion, our results show a phenotypic and proteomic characterization of an unusual cellular state of a replication-arrested strain generated by specifically targeting DnaA boxes 6 and 7 with CRISPRi, which disintegrates the replisome and in the inhibition of critical cell cycle processes like DNA replication and cell division, while protein translation and cell elongation persists.

## Materials and Methods

### Strains and culture conditions

Bacterial strains and vectors used in this study are listed in **Tables S1 and S2**. *B. subtilis* and its derivative strains were routinely cultured in Luria-Bertani (LB Broth Miller, Becton Dickinson) medium containing: 10 g/L tryptone, 5 g/L yeast extract, 10 g/L NaCl (and 15 g/L agar for solid medium) with constant shaking of 180-200 rpm at 37°C. In addition, LB was supplemented with glucose (1%) or xylose (1%) when indicated. *E. coli* strains were grown at 37°C with constant shaking at 180 rpm in LB or LB agar supplemented with the appropriate antibiotics for selection on plates.

*B. subtilis* was grown at 37°C on a LB agar plate streaked from bacteria glycerol stocks stored at -80°C. Subsequently, cultures were grown from single colonies. When needed, antibiotics were added to the media at the following final concentrations: 10 μg/ml kanamycin (CorningTM) and 1 μg/ml erythromycin (Sigma-Aldrich); 15 μg/ml lyncomycin (CorningTM) and 150 μg/ml spectinomycin (Sigma-Aldrich) for *E. coli*. Xylose was added to a final concentration of 1% (w/w) to induce the conditional promoter (P_*xylA*_). Glucose was used at a final concentration of 1% (w/v) and was used to repress the conditional inducible xylose inducible promoter (P_*xylA*_).

### Serial dilution plating viability assay

Overnight cultures were diluted to OD_600 nm_ 0.01 in 20 ml LB with 1% glucose in 100 ml flasks and further incubated at 37°C, 180 rpm until OD_600 nm_ 0.25 (+/- 0.025). To ensure cells are in early exponential phase, cultures were then back-diluted 1:10 in a total volume of 120 ml LB and 1% glucose in 1 L flasks, incubated at 37°C, 180 rpm until OD_600 nm_ 0.1 (+/- 0.025). Cells were pelleted at 4000g, at room temperature for 5 min and resuspended in LB supplemented with either 1% glucose or 1% xylose. Cells were incubated at 37°C, 180 rpm and 100 μl samples were taken at 0, 1, 3 and 20 h and then 10-fold serially diluted in LB. 5 μl of each dilution was spotted onto LB agar plates containing 1% (w/v) glucose to inhibit additional expression of dCas9 and incubated overnight at 37°C. The number of CFU/ml was monitored.

### Determination of origin-to-terminus ratio by qPCR

Cells in early logarithmic phase (OD_600 nm_ 0.1 ± 0.025) were pelleted at 4000 *g*, at room temperature for 5 min, split in two and resuspended either in LB supplemented with 1% glucose or 1% xylose and incubated at 37°C, 180 rpm. 15 ml samples were taken at different time points (0, 3 and 20 hours), spun down at 11000 *g*, 4°C for 5 min followed by a genomic DNA extraction with the Nucleospin DNA extraction Kit (macherey-nagel) according to the manufacturer’s instructions.

The primer pair targeting the origin region was OLEC11491 and OLEC11492, and for the terminus was OLEC11493 and OLEC11494. qPCR reactions of 20 μl contained 2.5 ng of DNA, 200 nM of each primer and 10 μl of 2X Power SYBR™ Green PCR-Master-Mix (Applied Biosciences™), and amplifications were performed on a QuantStudio™ 5 Real-Time PCR system (Applied Biosciences) according to the following protocol: 95°C for 3 min, followed by 40 cycles of 95°C for 30 s, 60°C for 30 s and 72°C for 30 s. ori-ter ratios were analyzed using the 2ΔΔCt method(50). A fixed sample of the WT strain grown into late stationary phase, where the population would be expected to have an ori-ter ratio corresponding to 1, was used for normalization in every cycling run.

### Replisome localization by GFP-DnaN

Cell in early logarithmic phase were pelleted at 4000 g, at room temperature for 5 min and resuspended in LB supplemented with either 1% glucose or 1% xylose. Afterwards, they were incubated at 37°C, 180 rpm and 200 μl samples from timepoint 1, 2, 3 and 5 h were spun down at RT, 4000 g for 5 min. Cell pellets were washed twice with 1X PBS and resuspended in a final volume of 500 μl 1X PBS from which 1.5 μl were spotted onto 1.5% agarose pads and observed under the microscope using the GFP channel.

### DNA damage response assay

The promoter of *yneA* was fused to *gfp* gene and integrated into the ectopic *amyE* locus (27). Strains in early logarithmic growth phase (OD_600 nm_ 0.1) were treated with 1% glucose to repress or 1% xylose to induce expression of dCas9 for 3 hours. Cells treated with 3 μg/ml mitomycin for 3 hours were used as positive controls. 1 ml of each culture was washed twice (4000 *g*, room temperature for 5 min) and suspended in 1X PBS (phosphate-buffered saline, NaCl 137 mM, KCl 27 mM, Na_2_HPO4 10 mM, KH_2_PO_4_ 1.8 mM, pH 7.4); 2 μl were placed on 1.5% agarose in TAE pads.

### Fluorescence microscopy

Cell were grown to an OD_600 nm_ of ∼0.4. One microliter of cells was spotted onto 1.5% agarose pads and imaged. Images were acquired with an Inverted Microscope (Leica DMi8, DFC9000 GT VSC-D6212 camera), and a 100× phase contrast objective (HC LP APO 100X/1.40 oil). Filter sets for GFP channel were used when indicated.

### Time-lapse microscopy

After preparation of the culture, bacterial samples were added to an 1% agarose pad in S7_50_ media (51) supplemented with IPTG (1mM) and 1% xylose. Fluorescence time-lapse microscopy was carried out using a Ti-2 Nikon inverted microscope equipped with a CFI Plan Apochromat DM 60× Lambda oil Ph3/1.40 objective (Nikon) and camera Fusion FT (12bit sensitive scan mode standard, Hamamatsu) in the absence of binning. A denoise module (Nikon) was used during the acquisition of the images. Microscope settings were set to the following: GFP 488 nm 50ms 5%, Phase Contrast: 150ms DIA 12bit sensitive single plane, using a LED Lamp Lumencor Spectra III light engine (Lumencor) and filter cube MXR00256 - LED-DA/FI/TR/Cy5/Cy7-A (DAPI / FITC / TRITC / Cy5 / Cy7 - Full Multiband Penta) (Semrock). Images were taken every 10 minutes in a pre-warmed chamber at 37°C with Nikon Perfect Focus Systems (PFS4).

Image analysis was performed using Fiji (52) and ilastik (53) to segment the cells, and MicrobeJ (54) to subsequently detect the cells and analyse the mean GFP fluorescence intensities and cell length. For each strain, three positions on the agarose pad were imaged and analysed.

### Identification and quantification of proteins by mass spectrometry

#### Sample collection

Strains were grown in LB and 1% glucose. Once they reach early logarithmic phase cells pelleted at 4000 *g*, at room temperature for 5 min and resuspended in LB supplemented with 1% xylose. Cells were incubated at 37°C, 180 rpm and 20 ml samples were taken at different timepoints per treatment (0, 3, 4, 5 and 6 hours) followed by centrifugation at 11,000 *g*, 4°C for 5 min. Pellets were resuspended in 20 ml of ice-cold 1X PBS.

Cells were lysed with 500 µl 2x lysis buffer (final concentration: 2% SDS, 20 mM TCEP, 80 mM CAA (Chloroacetamide, Sigma-Aldrich), 100 mM HEPES pH 8 (VWR), 0.5x protease inhibitors (Roche) in tubes containing 25 mg 0.1mm silica beads (BeadBeater® Glass beads, 0.1 mm (Roth) and were disrupted using a FastPrep-24TM 5G Homogenizer (MP Biomedicals). Samples were spun down at 11000 g, 10 min to separate the soluble from the insoluble fraction. The supernatant was heated at 95°C for 5 min and then cooled down to room temperature. The protein concentration was determined using Pierce™ BCA Protein Assay Kit (ThermoFisher Scientific), followed by nucleic acid digestion with 0.5 Units of Benzonase per 1 µg of protein (Sigma-Aldrich) for 30 min at 37°C. Samples were kept at -20°C for further analyses.

#### Sample preparation

All samples were subjected to SP3 sample preparation(55) on an Agilent BRAVO liquid handling robot. Ten µg of a 1:1 mixture of hydrophilic and hydrophobic carboxyl-coated paramagnetic beads (SeraMag, #24152105050250 and #44152105050250, GE Healthcare) were added for each µg of protein. Protein binding was induced by addition of acetonitrile to a final concentration of 50% (v/v). Samples were incubated for 10 min at room temperature. The tubes were placed on a magnetic rack, and beads were allowed to settle for 3 min. The supernatant was discarded, and beads were rinsed three times with 200 µL of 80% ethanol without removing the tubes from the rack. Beads were resuspended in digestion buffer containing 50 mM triethylammonium bicarbonate and both Trypsin (Serva) and Lys-C (Wako) in a 1:50 enzyme to protein ratio. Protein digestion was carried out for 14 hours at 37°C in a PCR cycler. Afterwards the supernatant was recovered dried down in a vacuum concentrator.

#### Peptide labeling and fractionation

TMT 11plex (Pierce, #A37725) was used for peptide multiplexing and quantification. Briefly, equal amounts of peptides were resuspended in 50 mM HEPES pH 8.5. Additionally, 10% from each sample was pooled to create a common sample as internal standard. TMT reagents were allowed to equilibrate to room temperature for 30 minutes and were dissolved in anhydrous acetonitrile to a final concentration of 59 mM. To each sample TMT was added to a final concentration of 11.8 mM and tubes were incubated at 25°C for 60 minutes with mixing at 500 rpm on a ThermoMixer. Labeling was quenched by addition of hydroxylamine to a final concentration of 0.4%. Samples were mixed, desalted using solid phase extraction (Seppak 1cc/50mg, Waters), dried down in a vacuum concentrator and resuspended in 20 µL 2% acetonitrile. Basic reversed phase fractionation was performed on a quaternary Agilent 1290 Infinity II UPLC system equipped with a Kinetex Evo-C18 column (150 × 2.1 mm, 2.6µm, 100 Å, Phenomenex) that was operated at 40°C. Solvent A consisted of HPLC grade water, solvent B consisted of 100% acetonitrile, and solvent C consisted of 25 mM ammoniumbicarbonate in water. Fractionation was carried out at a constant flow rate of 100 µl/min using a linear gradient from 2-25% acetonitrile within 50 minutes, followed by column washing and equilibration. Over the whole gradient solvent C was kept constant at 10%. In total 32 fractions were collected in conical 96well plates. The organic solvent was removed in a vacuum concentrator for 30 minutes and fractions were concatenated into 8 final samples. Peptides were acidified with formic acid, desalted using OASIS HLB 96well cartridges (Waters, #186001828BA), dried down and resuspended in 2% acetonitrile, 0.1% trifluoroacetic acid (TFA) prior MS analysis.

#### Mass spectrometry

All samples were analyzed on a Orbitrap Exploris (Thermo Scientific) that was coupled to a 3000 RSLC nano UPLC (Thermo Scientific). Samples were loaded on a pepmap trap cartridge (300 µm i.d. x 5 mm, C18, Thermo) with 2% acetonitrile, 0.1% TFA at a flow rate of 20 µL/min. Peptides were separated over a 50 cm analytical column (Picofrit, 360 µm O.D., 75 µm I.D., 10 µm tip opening, non-coated, New Objective) that was packed in-house with Poroshell 120 EC-C18, 2.7 µm (Agilent). Solvent A consists of 0.1% formic acid in water. Elution was carried out at a constant flow rate of 250 nL/min using a 180 minute method: 8-33% solvent B (0.1% formic acid in 80% acetonitrile) within 120 minutes, 33-48% solvent B within 25 minutes, 48-98% buffer B within 1 minute, followed by column washing and equilibration. Data acquisition on the Orbitrap Exploris was carried out using a data-dependent method in positive ion mode. MS survey scans were acquired from 375-1500 m/z in profile mode at a resolution of 120,000. AGC target was set to 100% at a maximum injection time of 50 ms. The ten most abundant peptides were isolated within a 0.4 m/z window and subjected to HCD fragmentation at a normalized collision energy of 36%. The MS2 AGC target was set to 200%, allowing a maximum injection time of 54 ms. Product ions were detected in the Orbitrap at a resolution of 30,000. TurboTMT acquisition as enabled. Precursors were dynamically excluded for 45 s.

#### Data analysis

Data analysis performed as described in Schäfer and collaborators (56). A two-tailed t-test and *p*-value correction were performed using Perseus to identify differentially expressed proteins (|log_2_ fold change| ≥ 1; *p-*value ≤ 0.05). Enriched pathways, biological processes and Uniprot keywords were calculated using DAVID (57, 58).

Proteins forming part of the ribosome or involved in cell length, shape, or cell division used for profile plots and heat maps were retrieved from SubtiWiki (59). The log2 protein intensities were scaled to standard deviation units (*z*-scores) using R for profile plots.

#### Data availability

The mass spectrometry proteomics data have been deposited to the ProteomeXchange Consortium via the PRIDE (60) partner repository with the dataset identifier PXD036876.

## Supporting information

VideoS1-CRISPRi

## Acknowledgments

We acknowledge Prof. Heath Murray from Newcastle University for valuable discussions and for providing strains that facilitated the cloning of strains EC3237 and EC3266. In addition, we thank Tim Sullivan for support with sgRNA off-target scoring. Finally, the authors are grateful to the members of the Charpentier group for constructive discussions and critical reading of the paper. This work was supported by the Max Planck Society [to M.E. and E.C.], the Max Planck Foundation [E.C.], Deutsche Forschungsgemeinschaft (DFG) [Leibniz-Prize to E.C.] and the Volkswagen Stiftung [to A.K.W.E.].

## Contributions

V.M.G, A.K.W.E., K.T., F.A.C., C.K.F. and E.C. designed the study; V.M., K.S., C.K.F and M.H. performed the experiments; F.A.C., M.H., C.K.F. and V.M.G performed the data analysis; V.M.G., F.A.C., M.H. and A.K.W.E. interpreted the data; A.K.W.E., K.T., M.E., and E.C. oversaw the project; V.M. wrote the paper. All the authors read, edited, and approved the paper.

