## Supplementary table S1-S3, figures S1-S3, Legend video S1 for "*Bacillus subtilis* remains translationally active after CRISPRi-mediated replication initiation arrest"

### Supplementary information

#### Table S1. Bacterial strains used in this study

| Strain | Relevant characteristics | Source |
|---|---|---|
| ***Escherichia coli* strains – Hosts for cloning** | | |
| **TOP10** | K12 strain, $F^{-}$ *mcrA* Δ*(mrr-hsdRMS-mcrBC)* φ80d*lacZ*Δ*M15* Δ*lacX74 recA1 araD139* Δ*(ara-leu)7697 galU galK* $\lambda^{-}$ *rpsL endA1 nupG*; SmR | Invitrogen |
| **NEB 5-alpha** | K12 strain, *fhuA2*::IS2 Δ*(mmuP-mhpD)169* Δ*phoA8 glnX44* φ80d[Δ*lacZ58(M15)*] *rfbD1 gyrA96 luxS11 recA1 endA1* $rph^{WT}$ *thiE1 hsdR17* | NEB, Anton & Raleigh, 2016 (1) |
| ***Bacillus subtilis* strains** | | |
| **EC3017 (WT)** | PY79<br>wildtype, *trpC2* prototroph | Youngman et al, 1984 (2) |
| **EC3137 (WT-dCas9)** | EC3017 *lacA*::$P_{xyl}$-*dcas9* ($Erm^R$) | This work |
| **EC3146 (CRISPRi$^{box1-2}$)** | EC3137 *amyE*::$P_{veg}$-sgRNA$^{box1-2}$ ($Cm^R$, $Erm^R$) | This work |
| **EC3147 (CRISPRi$^{box6-7}$)** | EC3137 *amyE*::$P_{veg}$-sgRNA$^{box6-7}$ ($Cm^R$, $Erm^R$) | This work |
| **EC3149 (CRISPRi$^{box3-4}$)** | EC3137 *amyE*::$P_{veg}$-sgRNA$^{box3-4}$ ($Cm^R$, $Erm^R$) | This work |
| **EC3686 (CRISPRi$^{nc}$)** | EC3137 lacA::Pxyl-dCas9, amyE::Pveg-non-coding sgRNA ($Cm^R$, $Erm^R$) | This work |
| **EC3237** | EC3017 $P_{dnaAN}$-*dnaA-gfpdnaN* ($Cm^R$) | This work<br>Adapted from (3) |
| **EC3259** | EC3017 *amyE*::$P_{veg}$-sgRNA$^{box6-7}$, *lacA*::$P_{xyl}$-*dcas9*, *gfpdnaN* ($Spc^R$, $Erm^R$, $Cm^R$) | This work |
| **EC3266** | EC3017 *amyE*::$P_{yneA}$-*gfp-yneA* ($Cm^R$) | This work<br>Adapted from (4) |
| **EC3270** | EC3017 *amyE*:: $P_{yneA}$-*gfp-yneA, thrC*::$P_{veg}$-sgRNA$^{box6-7}$ ($Cm^R$, $Spc^R$) | This work |
| **EC3272** | EC3270 *lacA*::$P_{xyl}$-*dcas9*($Erm^R$) | This work |
| **EC3556** | EC3017 *thrC*::$P_{hyperspank}$-*gfpmut2* ($Spec^R$) | This work |
| **EC3673** | EC3556 *amyE*::$P_{veg}$-sgRNA$^{box6-7}$, *lacA*::$P_{xyl}$-*dcas9* ($Spc^R$, $Erm^R$, $Cm^R$) | This work |

**Abbreviations:** Antibiotic resistances are represented as: $Amp^R$ – ampicillin, $Cm^R$ – chloramphenicol, $Kan^R$ – kanamycin, $Spc^R$ – spectinomycin, $Erm^R$ – erythromycin.

**Table S2. Plasmids used in this study**

| Plasmid | Relevant characteristics[b] | Source |
|---|---|---|
| **pEC2728 (pJMP1)** | *B. subtilis lacA/ganA* integrative vector *pAX01ΩP$_{xyl}$-dcas9* (Erm$^R$) | Addgene #79873 (5) |
| **pEC2728 (pJMP2)** | *B. subtilis amyE* integrative vector *pDG1662ΩP$_{veg}$-sgRNA_RR1* (Cm$^R$) | Addgene #79874 (5) |
| **pEC2730 (pJMP3)** | *B. subtilis thrC* integrative vector *pDG1731* ΩP$_{veg}$-*sgRNA_RR1* (Spc$^R$) | Addgene #79875 (5) |
| **pEC2741** | *B. subtilis amyE* integrative vector *pDG1662ΩP$_{veg}$-sgRNA*$^{box1-2}$ (Cm$^R$) | This work |
| **pEC2742** | *B. subtilis amyE* integrative vector *pDG1662ΩP$_{veg}$-sgRNA*$^{box6-7}$ (Cm$^R$) | This work |
| **pEC2743** | *B. subtilis amyE* integrative vector *pDG1662ΩP$_{veg}$-sgRNA*$^{box3-4}$ (Cm$^R$) | This work |
| **pEC2892** | *B. subtilis thrC* integrative vector *pDG1731* ΩP$_{veg}$-*sgRNA*$^{box6-7}$ (Spc$^R$) | This work |
| **pEC2990** | pEC2730 *thrC*::P$_{hyperspank}$-*gfpmut2* (Spc$^R$) | This work |
| **pEC3048** | *B. subtilis amyE* integrative vector *pDG1662ΩP$_{veg}$-sgRNA_ncsgRNA* (Cm$^R$) | This work |

**Table S3. Oligonucleotides used for marker frequency analysis**

| Oligo | Sequence 5′-3′ | F/R | Purpose/ Target |
|---|---|---|---|
| **OLEC11491** | GATCAATCGGGGAAAGTGTG | F | qPCR *ori-ter* ratio |
| **OLEC11492** | GTAGGGCCTGTGGATTTGTG | R | qPCR *ori-ter* ratio |
| **OLEC11493** | TCCATATCCTCGCTCCTACG | F | qPCR *ori-ter* ratio |
| **OLEC11494** | ATTCTGCTGATGTGCAATGG | R | qPCR *ori-ter* ratio |

F/R - Forward/Reverse. All oligonucleotides were ordered from Sigma-Aldrich.

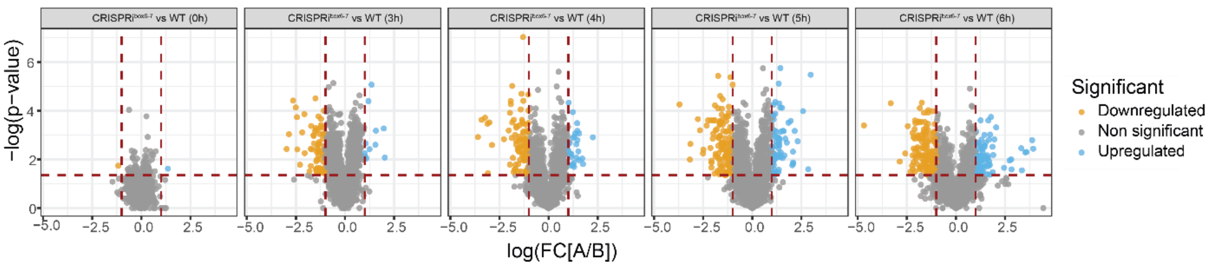


**Figure S1. *B. subtilis* proteomic response to replication arrest.**

Volcano plot of the WT strain versus the CRISPRi$^{box6\text{-}7}$ strain, showing the statistical significance (*p*-value) versus magnitude of change (fold of change), each protein is represented as a dot. Proteins with statistically significant differential expression (≥1.5-fold, *p*<0.05) located in the top right and left quadrants.

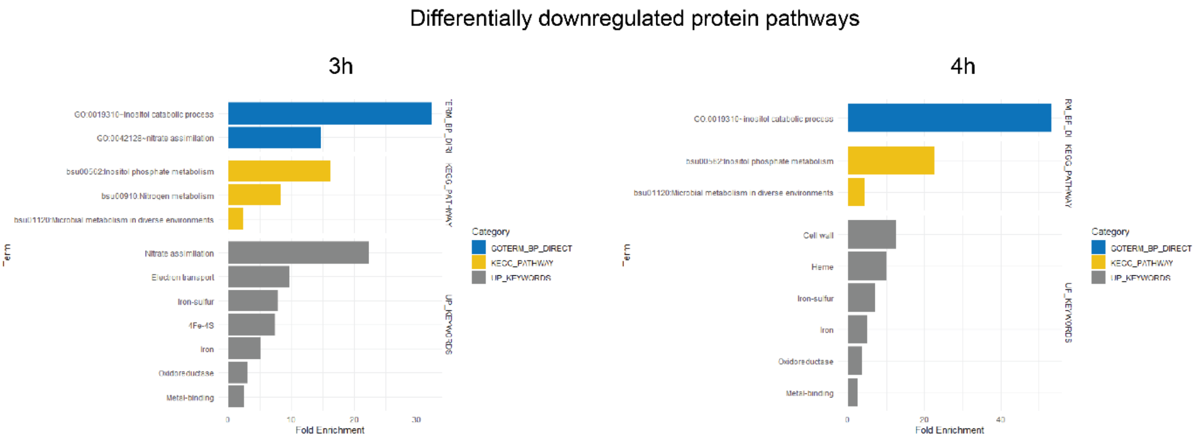


**Figure S2. Pathways showing depletion of proteins in CRISPRi$^{box6\text{-}7}$.**

The fold enrichment value represents the fraction of quantified proteins belonging to a particular category compared to the total number of proteins assigned to that category in the genome.

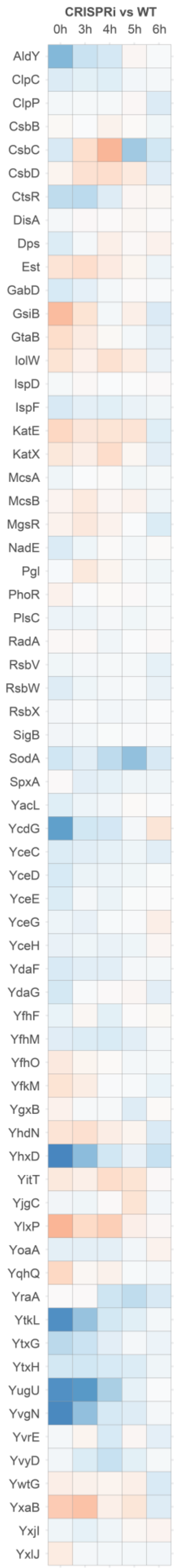


**Figure S3. No evidence of Sigma B mediated stress under replication arrest.**

Heatmap based on Fold Change values of proteins related to Sigma B regulon. All proteins annotations are retrieved from SubtiWiki.

**Video S1. Timelapse microscopy of WT and CRISPRi cells.**

Videos showing growth in $S7_{50}$ medium supplemented with IPTG and xylose.

WT and CRISPRi videos provided in an additional folder.
